# A highly scalable method for joint whole genome sequencing and gene expression profiling of single cells

**DOI:** 10.1101/2020.03.04.976530

**Authors:** Vasilios Zachariadis, Huaitao Cheng, Nathanael J Andrews, Martin Enge

## Abstract

Understanding how genetic variation alters gene expression - how genotype affects phenotype - is a central challenge in biology. To address this question in complex cell mixtures, we developed Direct Nuclear Tagmentation and RNA-sequencing (DNTR-seq), which enables whole genome and mRNA sequencing jointly in single cells. When applied to biobanked leukemia samples, DNTR-seq readily identified minor subclones within patients, as well as cell-type specific gene editing such as T-cell receptor rearrangements. mRNA-seq quality is equal to RNA-only methods, and the high yield combined with low positional bias of the genomic library preparation allows detection of sub-megabase aberrations at ultra low coverage of 0.5-3M read pairs per genome. Since each cell library is individually addressable, rare subpopulations can be re-sequenced at increased depth, allowing multi-tiered study designs where depth of sequencing is informed by previous results. In addition, the direct tagmentation protocol enables coverage-independent estimation of ploidy, which can be used to unambiguously identify cell singlets. Thus, DNTR-seq directly links each cell’s state to its corresponding genome at a scale enabling routine analysis of heterogeneous tumors and other complex tissues.

## Main

The identity of a cell is determined by the interplay between its epigenetic state and genome. In most healthy tissues of an organism, each cell’s genome is identical but due to differences in cellular concentration of transcription factors and other signaling molecules, its phenotype can be dramatically different. During cancer development, acquired genetic changes such as deletions, translocations and substitutions alters the cell state to promote growth. During disease progression, clones with additional genetic alterations will appear, leading to clonal heterogeneity followed by purifying selection and more aggressive disease. Single-cell whole genome sequencing (scWGS) allows us to take a snapshot of the clonal structure and heterogeneity in cell populations, and has been successfully applied to study tumors^1–4^ as well as naturally occurring genetic changes in healthy tissue^5–7^. However, this data is descriptive in nature and typically does not reveal the phenotypic traits of detected subpopulations.

Single-cell transcriptomics have proven to be a sensitive gauge of cell state and enabled researchers to characterize cell types from primary tissues of arbitrary complexity, as well as dissecting heterogeneity in gene regulation within these cell types. However, to determine the impact of genomic alterations on gene expression at single-cell resolution, we need to determine the full complement of transcript abundances and genetic variation in the same cell. Despite the promise of joint gDNA/mRNA-seq methodology, previous efforts^8,9^ have suffered from limitations related to technical biases, high cost or low specificity, and have not yet seen widespread adoption. On the other hand, microwell and droplet-based ultra-low coverage genomic sequencing methods have seen major methodological improvements recently, including the implementation of direct nuclear tagmentation strategies^10–13^. These DNA-only methods focus on optimizing library preparation throughput, enabling many cells from a single sample to be analyzed with the help of specialized equipment. For scientific questions where cells from many samples will be analyzed, however, the specialized and highly parallel nature of these methods can be a disadvantage.

We based DNTR-seq on a novel direct tagmentation protocol for genomic sequencing in an accessible multiwell format, combined with a sensitive full-length mRNA-seq protocol adapted from Smart-Seq2^14^. The protocol lends itself well to a 384-well based format which means that no specialized equipment is needed, while still allowing for routine analysis of many thousands of cells. Cells are deposited using FACS, adding surface protein expression as an optional layer of measurements for each cell, and allowing enrichment of rare populations by FACS gates where necessary. Our genomic library preparation is cost effective - at a typical ultra-low sequencing depth of 1M reads per genome, sequencing constitutes 80% of the cost per cell for scWGS. Importantly, individual cells can easily be re-sequenced to a higher depth allowing for flexible and iterative experimental designs.

## Results

### Development of an efficient protocol for joint mRNA and genome sequencing in single cells

In order to meet the requirements of low cost and low technical variability along with the throughput necessary for large scale genotype-phenotype experiments, we developed a method called Direct Nuclear Tagmentation and RNA-seq (DNTR-Seq), illustrated in Fig. 1a. We identified four main features that are necessary to make this method viable for widespread adoption: 1) High fidelity of both genomic and mRNA-seq. For genomic sequencing, this means avoiding lossy operations such as DNA cleanup as well as keeping PCR amplification cycles low, whereas for mRNA-seq we aimed for high sensitivity detection of full length transcripts. 2) Individually addressable cells, which is a requirement to allow sequencing cells at a variable depth - for example when performing a low-coverage sequencing pass and then hit-picking a subpopulation for deeper re-sequencing. 3) Easily adaptable by any genomics/scRNA-seq lab, no requirements on non-standard equipment. 4) Low gDNA positional bias. This facilitates identification of copy number variation (CNV) at ultra-low coverage, and means that less sequencing is required to meet a given minimum coverage.

**Fig. 1:**
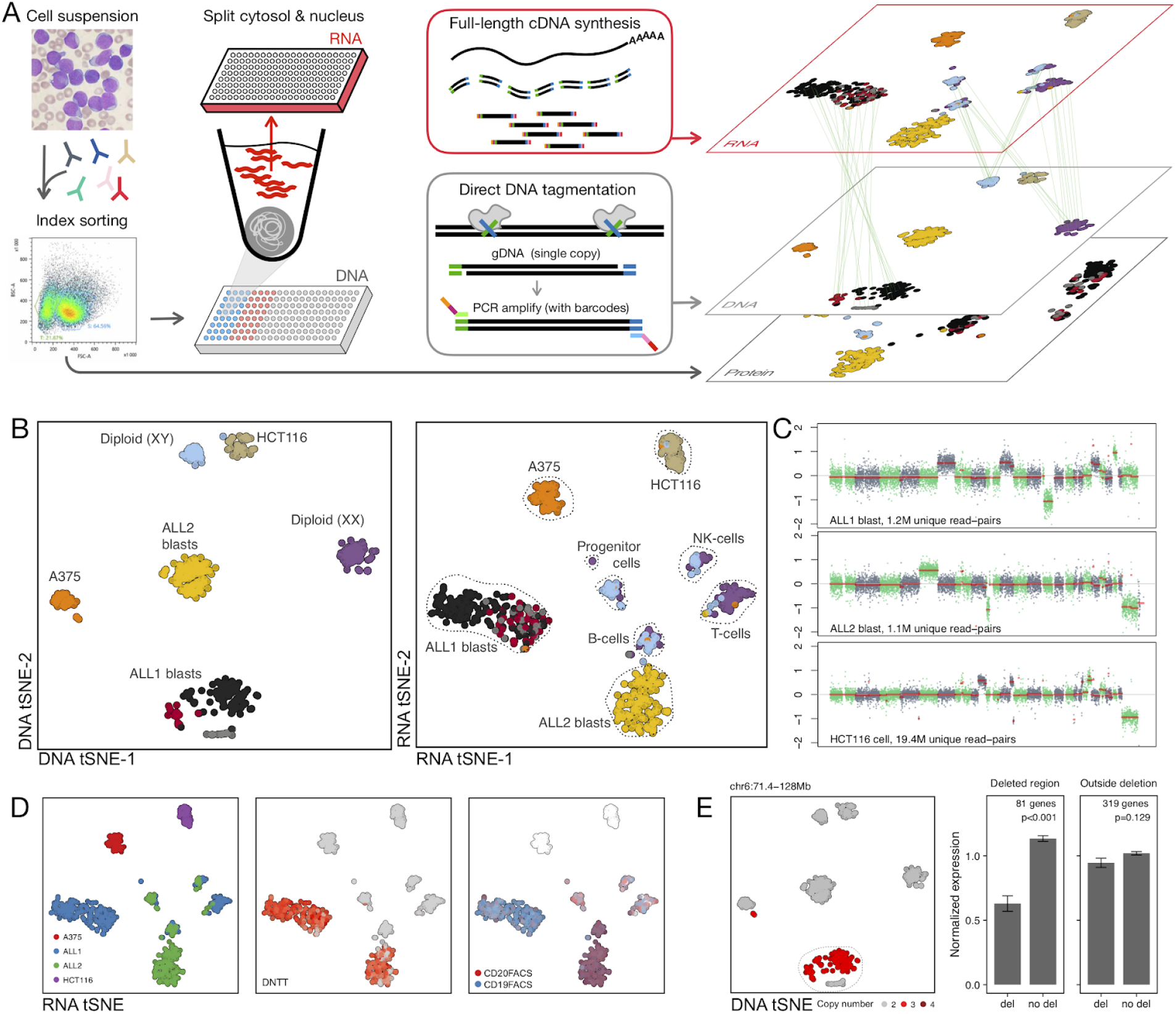
DNTR-seq allows joint profiling of single cells from patient samples or human cell lines. A) Schematic summary of the method. Right panel shows the same cells clustered by modality (mRNA, DNA CNV, cell surface protein expression), with colors indicating unsupervised classification based on CNV profiles. B) Analysis of CNVs based on DNTR-seq of 608 cells. Dimensionality reduction (tSNE) on joint copy number integers (left panel) and mRNA expression (right panel), colored by DNA class. C) Representative single-cell copy number profiles from ultra-low coverage sequencing (~1M read-pairs) of patient samples (ALL1, ALL2), and low coverage (~20M reads) sequencing of the cell line HCT116. D) Sample source, *DNTT* gene expression and cell surface protein expression (CD19 and CD20) overlayed on a tSNE of mRNA. E) tSNE of DNA, colored by copy number in a region of chr 6 (left panel), and normalized expression of genes in the affected region compared to non-affected (right panel). Deletion results in approximately dose-dependent reduction of mRNA abundance.

DNTR-seq is based on physical separation of the nucleus and cytosol into separate 384-well plates, followed by direct tagmentation of genomic DNA in the nuclear fraction and a template switching RNA-seq protocol on the cytosolic fraction^14^. Separation of the nuclear and cytosolic compartments is achieved by gentle lysis of the cell membrane followed by centrifugation and controlled aspiration, leaving the nucleus intact in the original sorting plate (Supplementary Fig. 1). The low cost of separation makes DNTR-seq ideal for large-scale experiments and allows flexible experimental designs. Since both fractions can be stored unprocessed for extended periods of time, we can for example detect rare cell populations first using scRNA-seq and analyze their gDNA further using genomic sequencing of selected wells. Finally, by avoiding pre-amplification and initial DNA purification, we achieve highly reliable ultra-low coverage DNA sequencing with low positional bias.

### DNTR-seq analysis of human patient samples and cell lines

We analyzed 608 cells using DNTR-seq from two pediatric acute lymphoblastic leukaemia (ALL) cases (ALL1 and ALL2), human colon adenocarcinoma cell line HCT116, and melanoma cell line A375 (Supplementary table 1). To demonstrate our ability to analyze biobanked material, we chose viably frozen patient samples which had been in cryogenic storage for 6 (ALL1) and 18 (ALL2) years. Regions with altered copy numbers were detected by segmenting gc- and mappability-corrected binned counts, and converted to absolute copy numbers by an expectation-maximization optimization procedure (see *Methods* for details). Unsupervised classification of absolute copy numbers yielded distinct clusters representing the variant genomes, with each cell line and leukemic blast clones separating into their own clusters. Normal diploid cells from ALL patient samples clustered according to XX and XY genomes, as expected (Fig. 1b, left panel). Copy number profiles were robust at the single-cell level, even with ~1M read pairs (Fig. 1c). In the joint transcriptional data, ALL1 and ALL2 blasts formed separate clusters by patient whereas normal B-cells, T-cells and macrophages instead clustered by cell type (Fig. 1b,d). ALL1 blasts harbored a subclonal partial loss of chromosome 6, with a dose-dependent decrease in expression of genes in the affected region (Fig. 1e). We also analyzed the TCR locus in T-cells, as classified by their transcriptional profile. Individual healthy T-cells all had unique TCR deletions, healthy non-T cells lacked deletions, whereas blasts were strictly clonal, with identical deletions, as expected (Supplementary Fig. 2). Together, these experiments highlight the complementary features of the two modalities, with gDNA- and mRNA-assays each showing specificity in classification, consistent with high-quality data. Joint analysis demonstrates that, as expected, tumorigenic alterations have a large impact on gene expression, whereas natural X/Y chromosome differences are largely silent.

### Direct nuclear tagmentation provides libraries with even coverage suitable for ultra-low pass sequencing

For whole-genome sequencing of the separated single nuclei, we developed a single-well protocol based on direct tagmentation using the hyperactive tn5 transposon system^15^. After disintegrating all nuclear protein by freezing and protease digestion, the free genomic DNA is digested by a transposome loaded with oligos compatible with barcoded Illumina sequencing adapters, which are subsequently added to the tagmented gDNA by PCR amplification. This three-step procedure is done without intermediate DNA clean-up, is highly efficient and is easy to automate. To benchmark the single-cell whole-genome sequencing (scWGS) element of DNTR-seq, we analyzed the ultra-low coverage sequencing data from ALL patients and cancer cell lines, including a few cells re-sequenced at higher coverage. At ultra-low coverages of 1-5M read-pairs/cell, duplicate rates were generally below 10%, increasing to 20.2%, on average, at a higher read depth (10-33M read-pairs; Fig. 2a and Supplementary Fig. 3). Since the DNA is tagmented prior to amplification, each genomic site can only be sequenced once per chromosomal copy, which allows us to calculate the yield based directly on library size. The duplicate rate indicates a total library size of 2.48E9 bp on average^16^, equivalent to a yield of 45.9% based on the mappable size of a diploid genome.

**Fig. 2:**
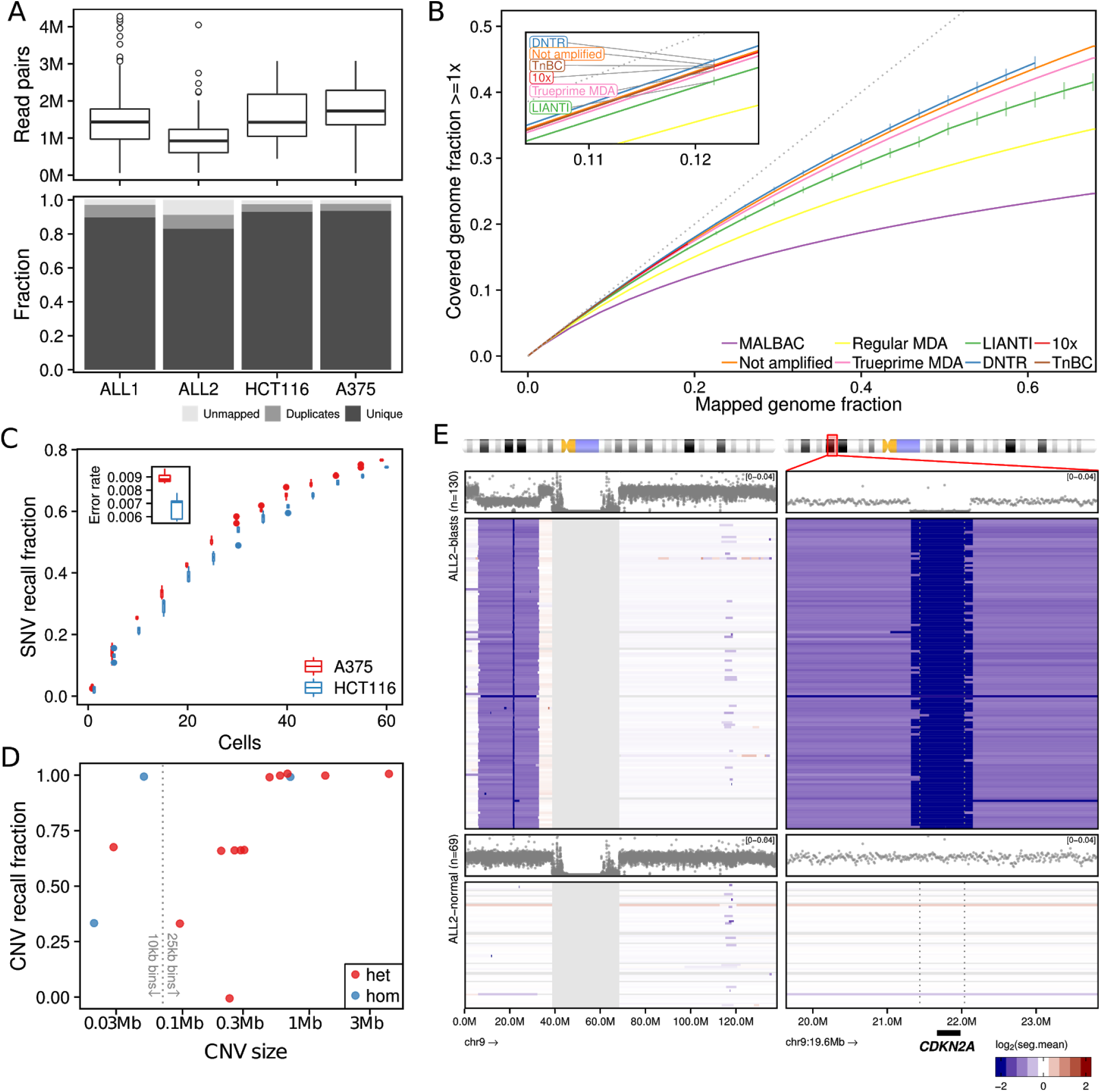
Ultra-low coverage sequencing using DNTR-seq produces libraries with high on-target rate and low positional bias. A) Sequencing characteristics of ultra-low coverage genomic DNTR-seq libraries from cell lines (A375, HCT116) and patient samples (ALL1, ALL2). B) DNTR-seq has low positional bias, comparable to unamplified bulk sequencing protocols. Lines show covered genome fraction as a function of the amount of DNA sequenced. Dotted line indicates the theoretical ideal condition x=y, when every new fragment sequenced provides new information, so that approaching this line indicates a more even distribution of DNA fragments. Where multiple replicates were available, vertical lines indicate ±1SD from mean. C) SNV detection using DNTR-seq. Shown is the probability of observing the correct allele with increasing number of sequenced cells. Inset indicates error rate, the probability of observing the incorrect allele at a homozygous position. D) Recall of CNV determined individually in single HCT116 cells sequenced to a depth of ~20M read pairs. E) Copy number calls of a *CDKN2A*/P16 locus deletion in ALL2 blasts. Copy number profile of chromosome 9 (left) and an expanded view of the sub-megabase deletion of CDKN2A (right panel, locus 9p21.3 highlighted in red in the ideogram above). Dotplot indicates aggregate copy number profile of blasts or normal cells, whereas individual copy number calls are shown as color-coded lines below each aggregate profile. Both a sub-megabase homozygous deletion and the surrounding heterozygous deletion is called in each leukemic blast cell, while absent in normal cells. Cell types (blast/normal) were determined by classification of the joint scRNA-seq data (Fig. 1b).

An important metric when comparing genomic sequencing methods is how evenly the reads are distributed in the genome. Positional bias can be caused, for example, by minute efficiency differences in exponential methods such as PCR or by sequence-dependent steps such as random priming^17^. Since the main cost of current scWGS methods is sequencing, considerable effort has been spent to minimize such biases. The direct tagmentation approach of DNTR-seq results in libraries with much more even coverage and with lower GC bias than traditional random priming methods such as MALBAC and MDA, and is equivalent to amplification-free bulk sequencing (Fig. 3b, Supplementary Fig. 4).

**Fig. 3:**
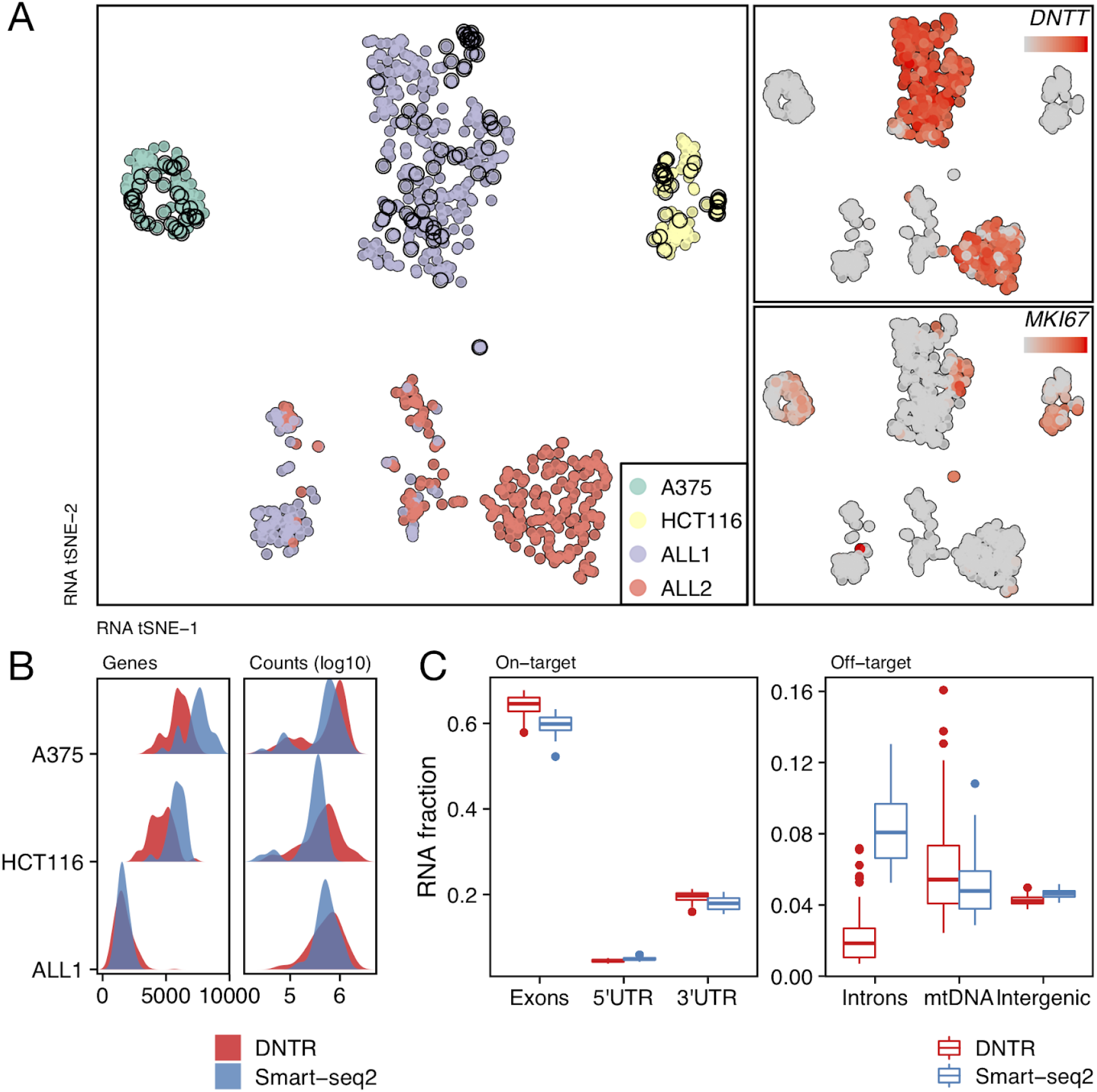
DNTR-seq mRNA libraries are comparable to Smart-Seq2 using whole-cell extracts. A) tSNE of mRNA libraries from single cells prepared using Smart-Seq2 or DNTR-seq analyzed in tandem. Libraries prepared using regular Smart-Seq2 (open black circles) are distributed throughout the correct clusters, indicating low bias between methods. B) Number of genes detected (left panel) and number of reads sequenced (right panel), in DNTR-seq and Smart-Seq2 libraries prepared under identical conditions. C) Fraction of mRNA-seq library reads mapping to gene features in HCT116 cells (Smart-Seq2 n=28, DNTR-seq n=25). On target reads from spliced transcripts are enriched in DNTR-seq libraries, whereas intronic and nuclear reads are enriched in regular Smart-Seq2 libraries.

Individual cells analyzed at ultra-low coverage will not reliably call single nucleotide variants (SNVs). However, by analyzing groups of clonal cells we can boost this capability - in our data, the recall probability of SNVs when using 60 single cells was around 80% (Fig. 2c). Importantly, variants detected in more than one cell implies strong error correction, since they must originate from different starting DNA template and are therefore unlikely to stem from early-round PCR errors.

The low positional bias of DNTR-seq makes it ideally suited to call copy number variation (CNV) within single cells. Ultra-low coverage sequencing is usually analyzed in a two-step procedure, where a rough copy number profile called on each cell in isolation is first used to determine clusters of clonal cells, then analyzed together to obtain refined breakpoints^11^. To test DNTR-seq’s single cell performance, we instead applied a method of copy number calling which considers each single cell on its own, which resulted in high recall of known sub-megabase CNVs (Fig. 2d,e). As shown in Fig. 2e, we could readily detect a sub-megabase *CDKN2A* deletion in every cell identified as an ALL blast using the joint transcriptional profile, whereas cells identified as non-leukemic did not carry the deletion.

### Transcriptome data from DNTR-seq has similar quality to Smart-Seq2 with a higher on-target fraction and less unspliced transcripts

We based the mRNA sequencing of DNTR-seq directly on Smart-Seq2^14^, the current gold-standard method for high-sensitivity full-length scRNA-seq. This allows detection of low-abundant transcripts, quantification of allele-specific expression and SNV calling on the mRNA-seq data. There are subtle differences between DNTR-seq and Smart-Seq2 owing to the exclusion of the cell nucleus in DNTR-seq: nuclear transcripts are expected to be largely absent from DNTR-seq data, and some cytosolic mRNA could be lost in the separation step, leading to loss of sensitivity. To directly compare DNTR-seq to Smart-Seq2, we prepared libraries from the human tumor cell lines A375 and HCT116, as well as one ALL patient sample, using both methods in parallel. In a standard dimensionality reduction analysis, the cells did not cluster as separate entities, and the Smart-Seq2 cells were well distributed within the correct cell clusters (Fig. 3a), indicating little or no impact on high-level transcriptome analysis (Supplementary Fig. 5). In patient samples, total read counts and number of detected genes were similar using the two methods, whereas the number of genes detected was slightly lower in the cell lines analyzed using DNTR-seq (Fig. 3a). Intronic regions, which are found in unspliced transcripts inside the nucleus, were less than half as common in DNTR-seq data compared to Smart-Seq2 data, underscoring the large differential effect on nuclear transcripts (Fig. 3c). Exonic reads, on the other hand, were relatively more highly abundant in DNTR-seq data, due to a lower fraction of off-target reads from intronic or non-genic regions. Taken together, this data shows that mRNA libraries from DNTR-seq are of similar quality to those from RNA-only methods, and can be used interchangeably in downstream analysis.

### DNTR-seq overlap analysis provides coverage-independent determination of ploidy or cell number

Single-cell analysis has proven particularly useful as a tool to find new cell types or intermediate cell states and reconstruct differentiation trajectories. However, we currently lack tools to unambiguously prove that a transcriptional profile originates from a single cell. Doublets of cells from two different cell types appear as intermediate transcriptional profiles and cannot be distinguished from a “true” intermediate cell. The direct tagmentation procedure of DNTR-seq, however, offers an orthogonal type of data that can directly determine whether the genetic material corresponds to a single genome (2n) or two genomes (4n). After direct tagmentation, the number of overlapping fragments at a specific genomic site (its *physical genomic coverage*) is restricted by the ploidy at that site. This is because, as shown schematically in Fig. 4a, each chromosome is directly cut into non-overlapping fragments by the transposome with no prior amplification. For a diploid (2n) cell, this means that a coverage of more than two is not possible, whereas for a tetraploid cell or cell doublet (4n) the maximum coverage is four. When using ultra-low coverage sequencing, the probability of observing >2x coverage is exceedingly low even for 4n genomes found in doublets. However, the difference in scarcity of template between a diploid cell compared to a tetraploid also affects the probability of observing 1x coverage compared to 2x coverage in a predictable manner (Fig. 4b). We can use this to calculate standardized coverage probabilities (SCP), where SCP[*k*] is the probability of a genomic position being covered by *k* fragments at a standardized sequencing depth (see *Methods* for details). When applied to wells where we intentionally deposited two cells, or cells containing multiple genomes (eg. S-phase/M-phase), the coverage probabilities unambiguously identified them as 4n (Fig. 4c).

**Fig. 4:**
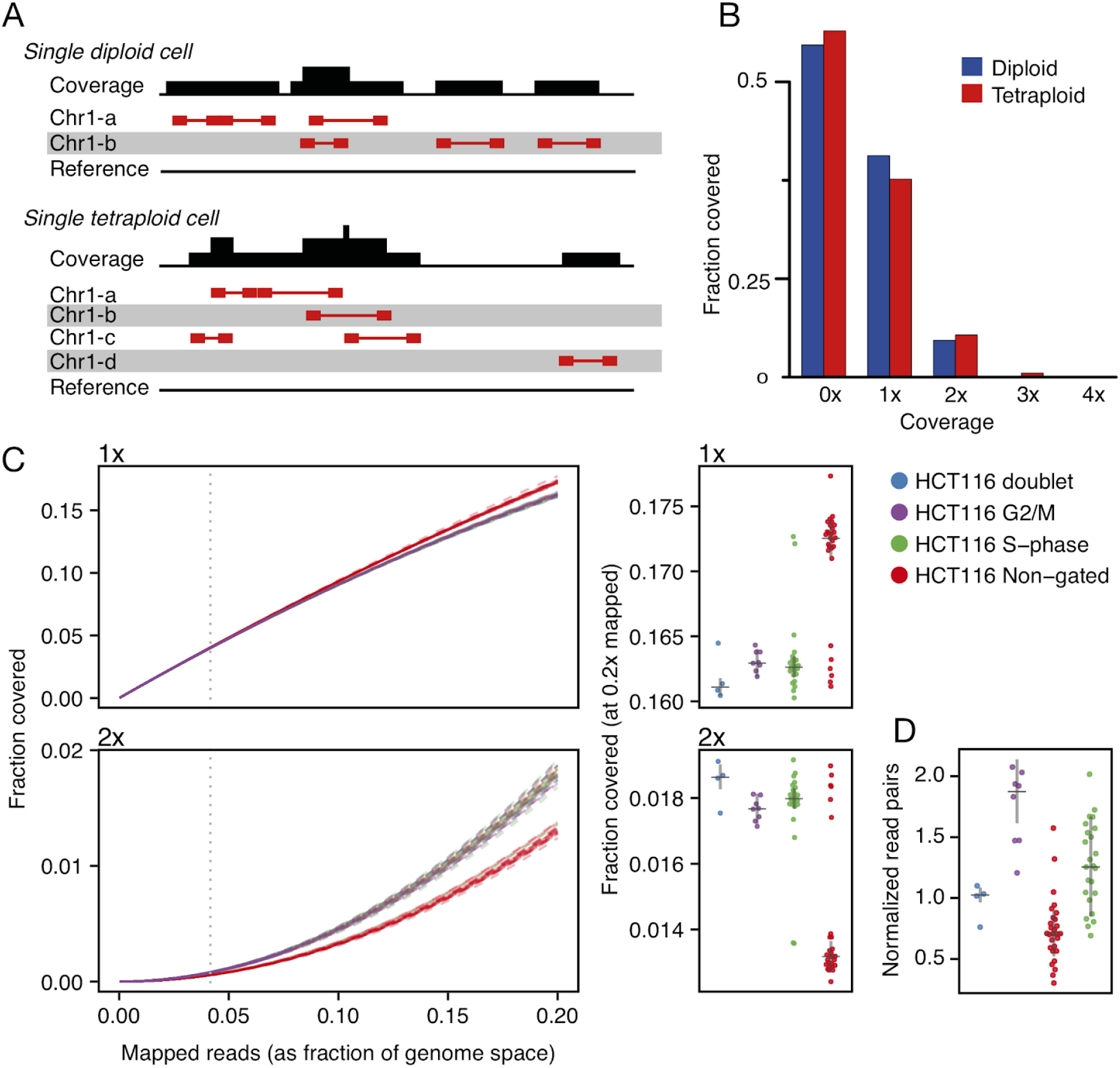
Standardized coverage probabilities (SCP) for genomic reads in single cells allows coverage-independent detection of doublets. A) Principle of overlap profiles. Paired-end reads are indicated by red connected boxes, aligned to the homologous template of origin. Top panel shows aligned reads to a single diploid genome (two sister chromosomes: Chr1-a and Chr1-b), and the bottom panel a tetraploid genome such as a cell doublet (Chr1a to d). A random distribution of tagmentation events will produce different coverage profiles in diploid and tetraploid cells. B) Simulated tagmentation of diploid and tetraploid genomes at 0.5x depth of coverage. C) SCP identifies 4n cells and cell doublets. HCT116 cells in G2/M or S-phase were selected using Hoechst staining (‘G2/M’, ‘S-phase’), or sorted as a mix of cell cycle states without gating (‘Non-gated’). Line plots (left) show the fraction of bases covered 1x or 2x as a function of sequencing depth. Vertical dotted lines indicate mean actual mapped base-pairs. Right panels show SCP (standardized to 0.2x sequencing depth), grey line is median +/− median absolute deviation. D) Normalized DNA library sizes of cells in C. Although doublet libraries are on average bigger than singlet libraries, the overlap is far too great to be useful for doublet detection.

Importantly, the SCPs are independent of sequencing depth. In theory, genomic sequencing depth could also be used as a proxy for ploidy, since two diploid genomes would be expected to result in a library double the size of one diploid genome (similarly to how we call chromosome gains/losses within a single genome). Indeed, ploidy estimation for CNVs (Fig. 1,2) or for late/early regions in DNA replication (Supplementary Fig. 6d) work remarkably well because they quantify *relative* differences within a cell. However, due to differences in tagmentation efficiency and exponential drift induced by differing PCR efficiency between wells, comparisons between library sizes of different cells are too noisy to give a reliable result, even when normalized by batch (Fig. 4d). In contrast, SCP completely sidesteps such variability between wells.

## Discussion

Studies of how genetic elements influence cell behaviour - correlating phenotype to genotype - are foundational to our understanding of how organisms function both in health^18^ and disease^19,20^. Single-cell mRNA-seq methodology can identify subtly different cell states in complex mixtures, and adding a genetic readout enables additional questions to be answered across various scientific fields. Coupling mRNA-seq to genetic information can either be done by joint analysis of mRNA and genomic DNA, or via forward-genetic screens. Such genetic screens in single-cell format are already proving valuable by allowing highly multiplexed experiments of gene function, but are intrinsically limited to *in vitro* or engraftment studies^21–23^.

Methods to jointly determine the transcriptional and genomic profile of single cells have been developed previously^8,9^, but have not yet seen widespread use. This is at least partly due to problems intrinsic to the methods, such as low sensitivity, need for specialized equipment, and/or high cost prohibiting experiments large enough to make meaningful inferences. DNTR-seq solves many of the problems with existing methods. No specialized equipment is needed and library preparation is easily automated using any standard liquid handling instruments. Our scWGS library preparation protocol is a one-well procedure with three enzymatic steps, and no intermediate clean up. This means that scWGS cost is mainly dependent on sequencing depth - at a typical ultra-low coverage of ~1M reads per cell, 80% of the cost is sequencing (which increases to ~99% at 20M reads/cell). Furthermore, we show that DNTR-seq produces libraries with lower positional bias than typical DNA-only methods, achieving the same breadth of coverage with fewer reads. Thus, the unbiased coverage of DNTR-seq directly translates into lower cost.

The multiwell format confers several advantages. First, sample processing can be split into discrete asynchronous steps - early nuclear separation allows mRNA-seq to be performed independently of WGS, and additional plates can be stored after sorting for later processing. Second, it allows us to add FACS-based measurements via index sorting, which provides a sparse proteomic layer to the data. Third, since the wells are independently addressable, it is possible to adjust sequencing depth in an iterative manner based on previous results. In this paper, we perform ultra low-pass sequencing on a few hundred cells and then re-sequence five genomes to a higher depth. However, since the sequencing required to detect a genetic aberration is directly dependent on its size, multiplexing can be taken much further for certain scientific questions. For example, rare tumor cells with a specific chromosomal loss could be selected in an initial screening step using 100-fold lower sequencing depth, and subsequently re-analyzed for transcriptional and genetic aberrations at higher resolution.

In addition to providing libraries with low positional bias, a feature of direct tagmentation of a single DNA template is that it produces non-overlapping fragments separated by shared Tn5 cut sites (Fig. 4a). These constraints are not affected by biases in library construction and have several potential uses. In this paper we use the coverage patterns to infer ploidy by calculating a standardized overlap probability, and demonstrate its use in detecting cell doublets. In addition, the fact that one tagmentation event creates two directly adjacent reads can be used for haplotyping. For a single chromosome copy, the genomic coordinate of a Tn5 cut site represents a unique molecular identifier^24^, and the two reads it separates must originate from the same DNA molecule. At a high enough sequencing coverage, chaining together multiple adjacent fragments into larger blocks would enable haplotype phasing as part of the analysis. This feature is useful both for SNP phasing and to delineate complex rearrangements in heterogeneous tumor samples.

Coupling measurements of cell state and genetic variation in single cells promises to solve previously intractable problems, with wide-ranging applications across normal and disease biology, including non-malignant mosaicism and aging. Here, we demonstrate its use in investigating the transcriptional phenotype of minor clones in 2 decades old biobanked patient material, and in detecting cell doublets. However, DNTR-seq is broadly applicable to questions in which direct genotype-phenotype associations are required. We expect DNTR-seq to be an important tool in this emerging field, combining high-sensitivity measurements of cell state with unbiased and efficient DNA library preparation.

## Methods

### Collection and processing of patient samples

Diagnostic bone marrow samples from individuals with acute lymphoblastic leukemia had been collected and biobanked with prior guardian consent at the Childhood Oncology Unit at Karolinska University Hospital. At diagnosis, cells were Ficoll gradient-separated and cryopreserved in FCS with 10% DMSO in liquid nitrogen. In this study, cryopreserved cells were quickly thawed in 37°C DMEM medium, washed of remaining DMSO and removed of dead cells (Dead Cell Removal kit, Miltenyi Biotec) before antibody staining and cell sorting.

### Culture of human cell lines

HCT116 and A375 cells were cultured in DMEM medium supplemented with 10% FBS and 1% Penicillin/streptomycin at 37°C, 5% CO_2_ and humidified atmosphere. The cells were split and passed at 70-90% confluency. When harvesting cells for FACS sorting, cells were split and collected by 10 min centrifugation at 300g. The cell pellets were then resuspend in 1ml 1xPBS and passed through a 40-mm cell strainer.

### FACS sorting

ALL cells were Fc-blocked and stained with CD34 (BV421), CD10 (FITC), CD20 (PE), CD38 (APC), CD45 (AF700), and CD19 (PE-Cy7) according to the manufacturer’s protocol (Sony Biotechnology). After washing, cells were resuspended in 1xPBS with 5% FBS and kept on ice, protected from light. For cell lines, propidium iodide (0.5μg/ml with cells) was added to identify live/dead cells during FACS sorting. In S-phase experiments we added Hoechst 33342 (ThermoFischer, cat no H3570) to HCT116 and A375 cells, and gated for viable singlets, S-phase or G2/M-phase (Supplementary Fig. 6). Cells were sorted into plates with lysis buffer using a Sony SH800S, centrifuged at ~2000g for 5 min and snap-frozen on dry ice.

### DNTR-seq nuclear separation

Immediately after FACS of single cells into 384-well plates containing 3μl lysis buffer (H2O 1.965μl, RNase inhibitor 0.075μl, 10% Triton X-100 0.06μl, 10mM dNTP 0.75μl, 100μM dT 0.075μl, ERCC 1:1.2M dilution 0.075μl), the plates were placed at −80°C. Freezing ensures efficient lysis of the cell membrane, but leaves the nucleus intact. Cells can be kept at −80°C for several weeks without adverse effects to the data. Upon thawing, the intact nuclei in the plate were spun down at 500g for 5min, and immediately processed for nuclear separation. The separation step was performed in an Eppendorf EpMotion 5073m pipetting robot by transferring 2μl of lysis buffer from the sorted plate into an empty 384-well plate at low flow rate (2mm/s) and with an aspiration height of 0.9mm above the well bottom. The low flow rate together with the high aspiration height ensures that the nucleus is not accidentally transferred along with the cytosolic fraction. We have also successfully performed nuclear separation with the Hamilton STARlet liquid handling robot, the Gilson Platemaster semi-manual 96-well pipette (both with the same settings as the EpMotion), or a fully manual 8-channel pipette.

### DNTR-seq whole genome sequencing

The nuclear fraction was first treated with Proteinase K (2μl, 0.13mg/ml final concentration) for 60min at 50°C, to lyse the protein-rich nuclear membrane and free genomic DNA from chromatin, followed by heat inactivation for 30min at 80°C. Tagmentation was performed for 10min at 55°C, using 0.1μl Tn5 stock solution (2.8mg/ml purified psfTn5-c006, Addgene plasmid #79107, in 50% Glycerol) previously loaded with standard Illumina Tn5 adapters in tagmentation buffer: 1.6μl TAPS-PEG buffer (50mM TAPS, 25mM MgCl2, 40% PEG 8k. Final concentration 8% PEG, 5mM MgCl2, 10mM TAPS), 3.3μl H_2_O (final volume 5μl). Tn5 was inactivated by applying 2μl 0.2% SDS treatment and incubating for 10min at 55C. Barcoding PCR was done as a 20μl reaction in the same plate using PCR master mix: 3.2μl H_2_O, 4μl 5X master mix, 0.6μl dNTP, 0.4μl KAPA HiFi DNA Polymerase (Roche, cat no. KK2102), 0.2μl 10% Tween-20 and 1.6μl custom barcoded primers (3.75uM each, see Supplementary table 2 for sequence data). The PCR program was 72°C/3 min, 95°C/30 sec, [95°C/15 sec, 67°C/30 seconds, 72°C/1 min] × 18 cycles, 72°C/5 min, and then 10°C hold.

After PCR amplification, 1.5μl from each well was pooled into a single 1.5ml tube, and sequentially cleaned-up twice using SPRI-beads (at 0.9X volume). For ALL patient samples, specified wells based on joint mRNA analysis were picked individually, pooled and cleaned as described. Finished libraries were sequenced on Illumina NextSeq 550.

### DNTR-seq mRNA libraries

Full-length mRNA-sequencing of the transferred cytoplasmic fraction of the cells was based on the Smart-Seq2 protocol^14^. After primer annealing for 3 min at 72°C, 3μl of reverse transcription (RT) mix was added (0.475μl SmartScribe [Takara], 0.125μl RNase inhibitor [Takara, cat no. 2313A], 1μl 5X First strand buffer [Takara], 0.25μl of 100mM DTT, 1μl 5M Betaine, 0.03μl 1M MgCl2, 0.05μl 100μM TSO, and 0.07μl H_2_O). RT was run at 42°C for 90min and 70°C for 5min, with a 4°C hold. Next, 7.5μl cDNA pre-amplification mix was added (1.068μl of H_2_O, 6.25μl 2X Kapa HiFi HotStart ReadyMix, 0.125μl of 10μM IS_PCR primer, and 0.056μl of Lambda Exonuclease). The PCR program was 37°C/30 min, 95°C/3 min, [98°C/20 sec, 67°C/15 sec, 72°C/4 min] × 21 cycles, 72°C/5 min and 4°C hold. Post-PCR clean-up using 0.7X volume SPRI-beads was performed twice, using a magnetic 384-well rack. For tagmentation, 0.7μl of diluted cDNA (150pg/μl) was mixed with 1.55μl of tagmentation buffer and 0.25μl of Tn5 (as above) and incubated at 55°C for 10 min. Immediately upon completion, Tn5 was inactived by adding 1μl 0.2% SDS. Barcoding PCR was done in the same plate by adding 19.5μl PCR mix (13.25μl H_2_O, 5μl 5X master mix, 0.75μl dNTPs, and 0.5μl KAPA HiFi DNA Polymerase) and 2μl primers (as above). The total volume of 25μl was run at 72°C/3 min, 95°C/30 sec, [95°C/15 sec, 55°C/30 sec, 72°C/45 sec] × 12 cycles, and 10°C hold. Finally, 2.5μl/well was pooled and cleaned up with SPRI beads at 0.9X volume twice. Smart-Seq2 libraries in this paper were prepared with the same protocol as above. Finished libraries were sequenced on Illumina NextSeq 550.

### Genomic read alignment and copy number estimation

Genomic reads were trimmed for adapter sequences using TrimGalore and mapped to human genome reference build hg38 (with ERCC reads added). After duplicate removal (Picard MarkDuplicates), read-pairs with MAPQ <20 or reads mapping to centromeric regions, satellite repeats and regions with excess coverage (top 1% of 100bp mers) were filtered out. For each cell, filtered reads were summarized over variable sized bins based on equal genome mappability^25^. For copy number estimation, summarized bin counts at 250kb resolution were normalized to the mean bincount per cell, GC corrected, log_2_ transformed and segmented using DNAcopy^26^. Integer copy numbers for each segment were estimated from log_2_ segment means by particle swarm optimization of euclidean distances between actual values and predicted ploidies.

### DNA sequence statistics and comparisons

We compared DNA sequence coverage breadth (bases covered or spanned by at least one paired read) to other single-cell DNA protocols. We downloaded or extracted unprocessed FASTQ reads of external protocols, aligned and processed them in an identical fashion to DNTR libraries (TnBC: SRA accession PRJNA350295, 10X: https://www.10xgenomics.com/solutions/single-cell-cnv, LIANTI: SRA accession PRJNA379710, MALBAC, MDA and Trueprime MDA all from SRA accession number PRJNA341815). To account for different sequence read lengths, all libraries were converted to fragments (Read 1 + insert + read 2). Fragments of each cell were then randomly subsampled at increments of 50M covered bp to calculate the fraction of genome covered (or physical sequence coverage) as a function of base pairs sequenced and mapped.

For single nucleotide variants, SNP genotype data of HCT116 and A375 cell lines from Cosmic (https://cancer.sanger.ac.uk/cosmic) was used as a truth set. For each cell and position in the SNP truth set, reads supporting reference or variant bases were summarized using mpileup^2727^.

For copy number variants, deletions from Cosmic SNP-genotyping data were compared to DNTR libraries of HCT116 singlets at higher depth (n=3; mean 20M read-pairs/cell). Accounting for the lower specificity of CNV calls and their coordinates, as compared to SNVs, we filtered the truth set for deletions also present in a pseudo-bulk version of all HCT116 cells sequenced (n=57). Segmented DNA calls in each single cell was then compared to this truth set, requiring a 50% reciprocal overlap to be called a match.

### DNA absolute ploidy calculation

Aligned and de-duplicated DNA read-pairs were converted into fragments (R1 start to R2 end position) and trimmed 5bp from each end. Trimming ensures that any detected overlaps represent different molecules, in contrast to 9bp overlaps rendered from expected duplications at the tagmentation site of Tn5. Fragments were filtered to include only autosomal chromosomes, MAPQ >=20, insert size <1000bp and unique start positions for stringent de-duplication. Finally, fragments were subsampled at increments of 5000 and depth of coverage calculated. The fraction of the genome covered at 1x, 2x, and 3x is a function of the number of mapped reads and cell ploidy. This function is well described by polynomial regression with a linear and a quadratic term (eg. coverage ~ mapped_fraction + mapped_fraction^2^), with mean R^2^ of 0.988.

### RNA read alignment and processing

RNA sequence reads were trimmed of adapters and low quality bases using cutadapt^28^. Trimmed reads were aligned to human reference genome build hg38, with added ERCC reads, using STAR 2.5.2b^29^. After removal of duplicate reads using picard^30^, read counts were summarized over annotated genes (NCBI release 106) using HTSeq^31^. Cells with less than 20000 reads or low *ACTB* counts were filtered out. The cutoff for *ACTB* expression was set to the 0.01 quantile of the normal distribution with empirical mean and standard deviation. Remaining cells were processed using Seurat^32^, including log_2_ normalization, feature selection and scaling before Louvain classification and *t*-SNE for visualization.

## Data availability

Sequencing data is available on GEO (accession GSE144296).

## Code availability

All code to reproduce the data processing pipeline, including a graphical user interface for joint DNA/RNA-seq analysis, is available at the DNTR-seq github repository (https://github.com/EngeLab/DNTRseq).

