## Supplementary figures 1-6 for "A highly scalable method for joint whole genome sequencing and gene expression profiling of single cells"

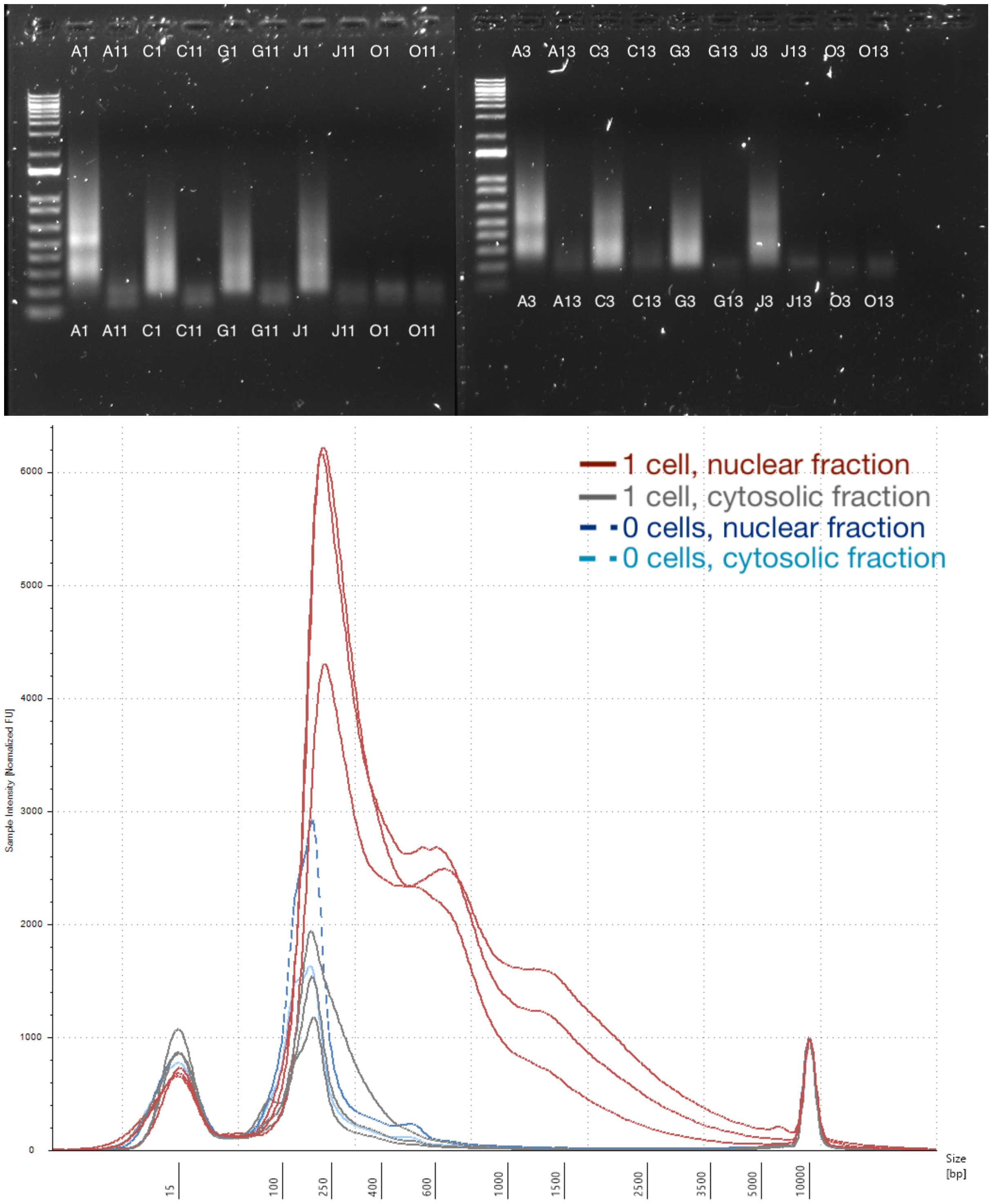


**Supplementary figure 1, Nuclear separation in DNTR-seq.** DNTR DNA-library preparation on paired nuclear and cytosolic fractions, after centrifugation and separation with liquid handling robot. *Top panel:* Starting with cells in 2µl (A1-O1) or 3µl (A3-O3) lysis buffer, 1.5 or 2µl, respectively, was transferred to corresponding wells (A3 → A13, C3 → C13 etc). A1/A3 are 20-cell positive controls, O1/O3 0-cell negative controls, the rest are single cells. Gel is a 1.5%

agarose gel with a 1kb+ ladder in the first well. *Bottom panel:* Size distribution of 3µl lysis transfers (C3/13 to O3/13 from top panel), with nuclear and cytosolic in red and grey respectively, and 0-cell controls in blue. Fragments were run on Agilent Tapestation DNA HS5kbp.

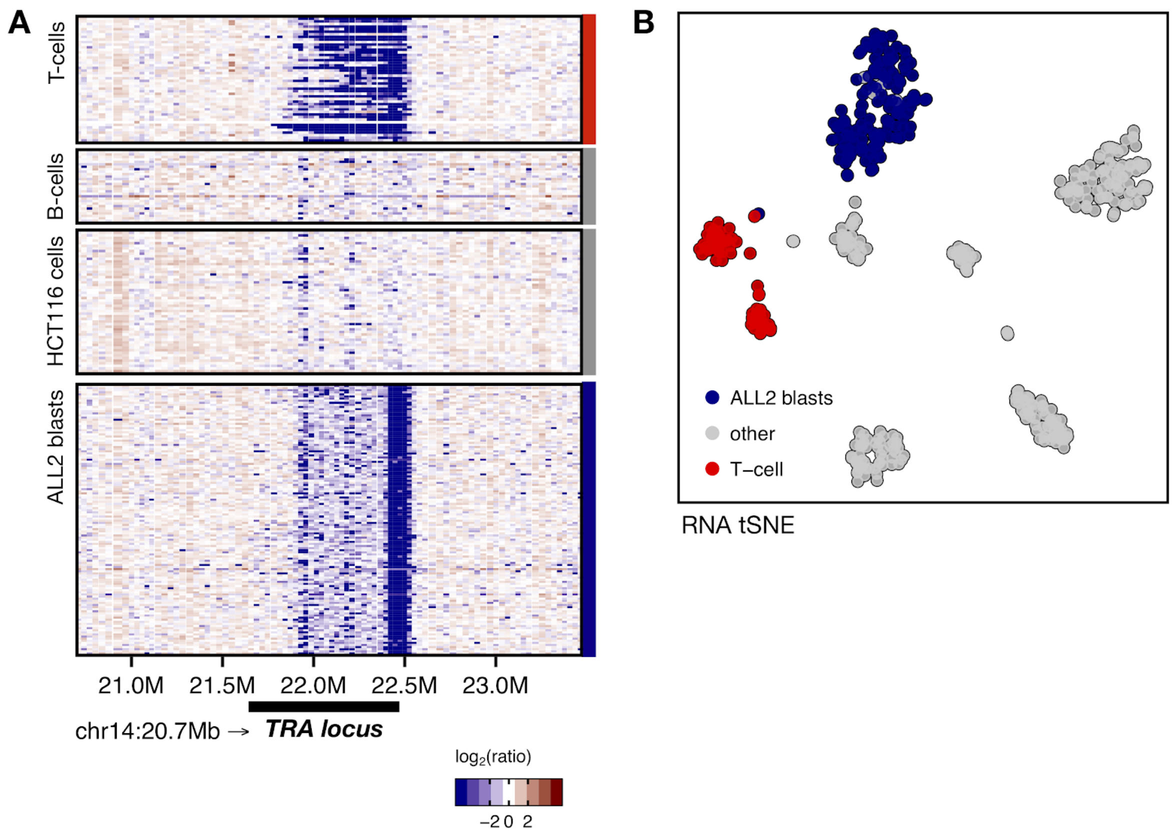


**Supplementary figure 2, DNTR-seq identifies individual TCR recombination events.** A) Per-bin ploidy in the T-cell receptor locus for cells identified based on their joint mRNA-sequencing data. T-cells (top) have varying TCR recombinations, whereas the ALL2 blasts (bottom) are clonal. B) tSNE of the cells in A.

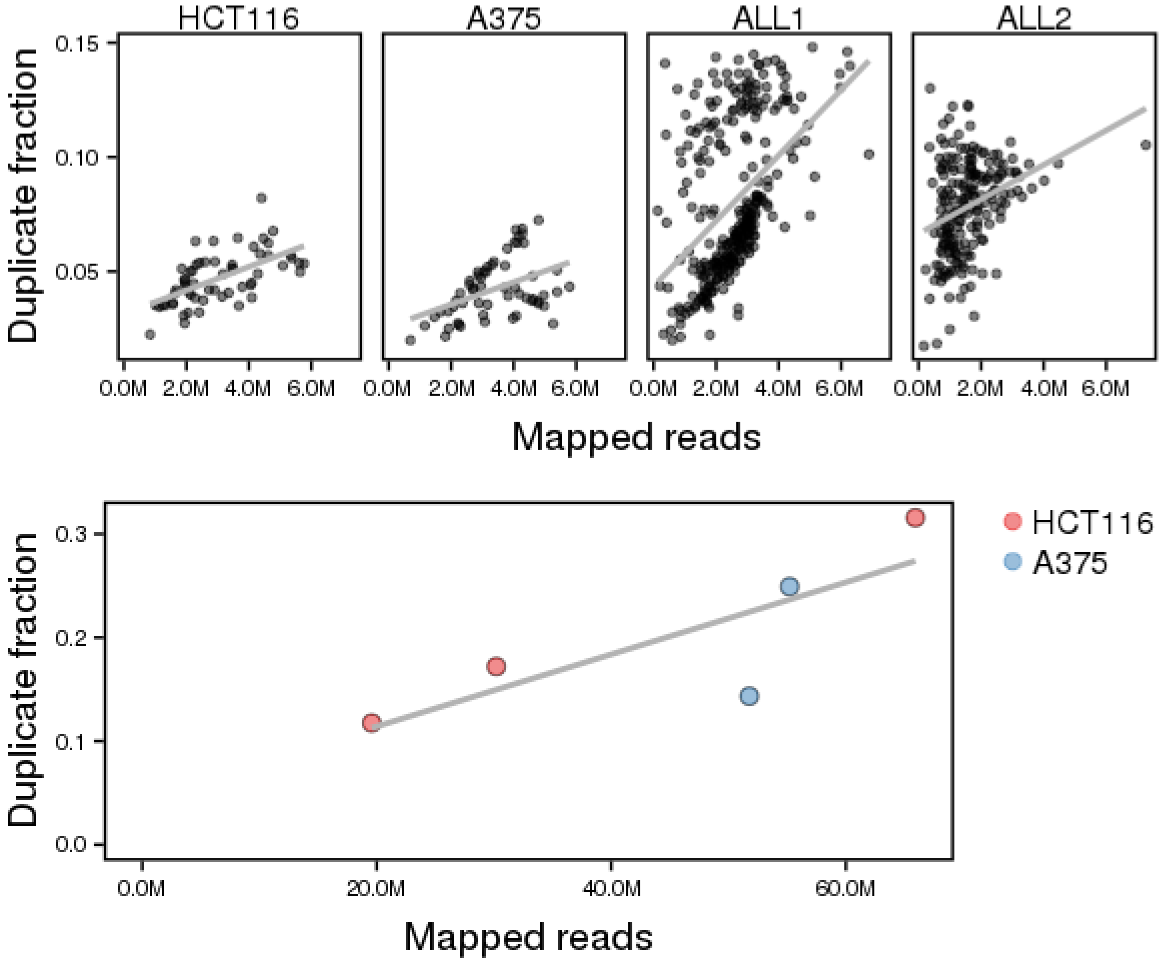


**Supplementary figure 3. Duplicate rate in WGS from DNTR-seq.** Total mapped reads versus duplicate fraction, in ultra-low (top, <5M read-pairs/cell) and low depth (bottom, 10-33M read-pairs/cell) sequenced single cells. Grey line indicates a linear model fit. ALL1 contains a subset of low-quality cells (as indicated by FACS analysis), with higher than normal duplication rates.

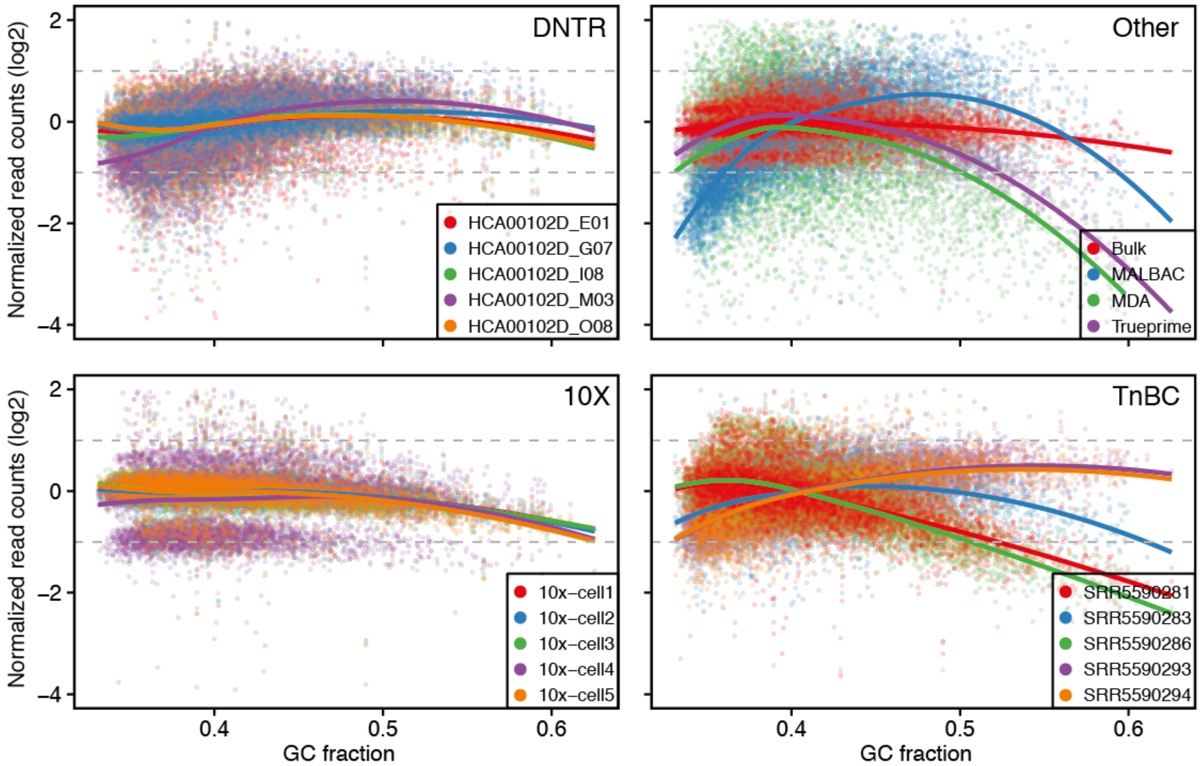


**Supplementary figure 4, comparison of GC bias in DNTR-seq WGS.** $Log_2$-transformed normalized read counts, summarized over 250kb bins, as a function of GC fraction. Panels are, clockwise, HCT116 single cells (DNTR-seq), compared to non-amplified bulk, MALBAC, MDA, Trueprime-MDA (all from PRJNA341815), 10X and TnBC (PRJNA350295). Colored lines are LOESS fit for each cell/library. Upper and lower dashed lines mark two-fold increase and decrease versus average observed coverage. Note that the DNTR-seq curve closely follows unamplified bulk.

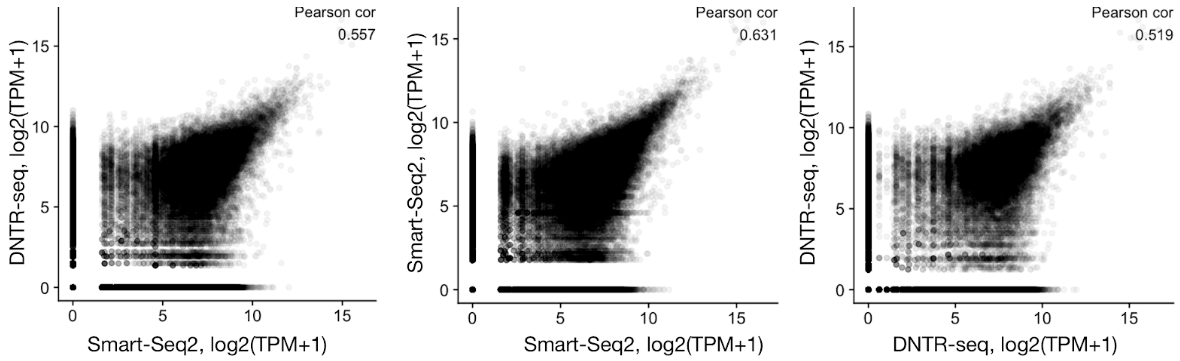


**Supplementary figure 5.** Correlation of mRNA expression, as $log_2$(TPM+1), between DNTR-seq and Smart-Seq2, as compared to within method correlation. Each panel shows expression of all genes from 10 x 10 individual cells.

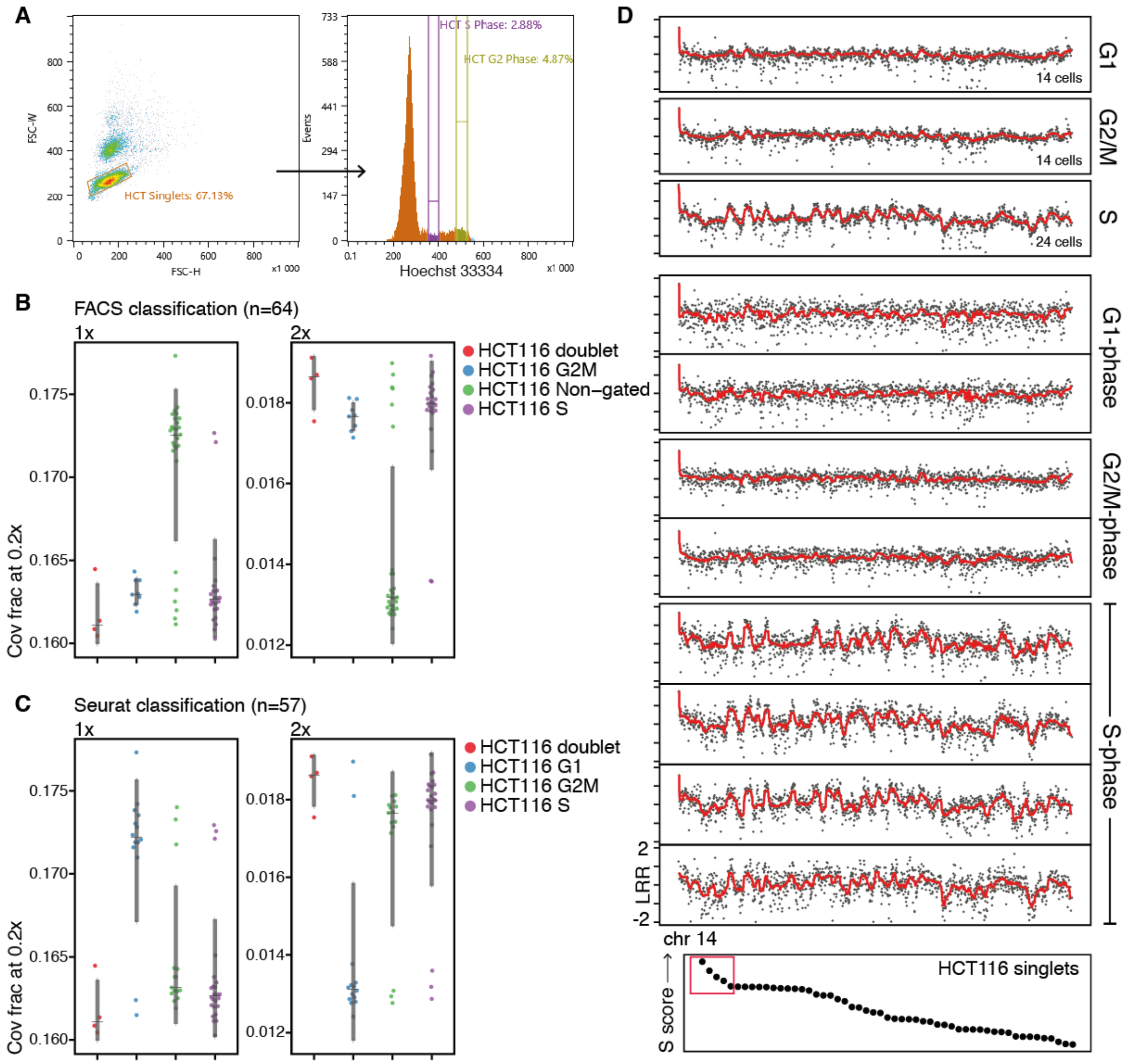


**Supplementary figure 6. Doublet detection and replication timing analysis using DNTR-seq.** A) Gating of HCT116 singlets, G2/M and S-phase cells. B) Standardized coverage probabilities (SCPs) colored by FACS or C) Seurat cell cycle scoring. D) S-phase replication timing in single cells. Top panels show means over all G1, G2/M and S-phase cells, as classified by Seurat. Bottom panels show single-cell DNA profiles from the different classes, G1 and G2/M are representative profiles, and S-phase profiles were selected based on S-phase score from Seurat (red box in bottom panel). Note that S-phase profiles show a distinct semi-conserved pattern of early to late replicating regions, while both G1 and G2/M profiles are flat. DNA profiles are $\log_2$-transformed normalized read counts over 50kb variable bins along chromosome 14. Red line indicates running median over 21 bins.
